# EcoHIV Treated Primary Microglia Exhibit a Highly Specialized and Sex-Dependent Transcriptional State

**DOI:** 10.64898/2026.10.25.761659

**Authors:** Brendan J. Veenstra, Kyler J. Nelson, Taylor R. Elder, Jill R. Turner, Kristen A. McLaurin

**Author notes:** **Address proofs and correspondence to:** Kristen A. McLaurin, Ph.D., Assistant Professor, Department of Pharmaceutical Sciences, College of Pharmacy, 789 South Limestone Street, University of Kentucky, Lexington, KY 40508.

## Abstract

The pathogenesis of HIV-1-associated neurocognitive disorders (HAND) is partly driven by microglial dysregulation. Indeed, HIV-1-induced microglial dyshomeostasis is characterized by a sustained pro-inflammatory state and exacerbated microglia proliferation. Herein, *in vitro* studies were conducted in primary microglia to elucidate the transcriptomic alterations underlying EcoHIV induced microglia hyperproliferation. Consistent with, and expanding upon, prior studies, treatment with EcoHIV exacerbated microglia proliferation evidenced by the upregulation of *Pcna* and *Mki67*. With regards to microglial receptors and adapter proteins, the transcriptomic expression of *Trem2* and *Dap12* was suppressed and increased, respectively, in microglia treated with EcoHIV. Crucially, dysregulation of the *Trem2/Dap12* signaling complex explained 51.4% of the variance in microglial proliferation in EcoHIV treated microglia. A preliminary investigation of the Trem2-mediated microglial survival and immunoregulation pathway affords additional support for a unique EcoHIV-induced transcriptomic profile. Specifically, independent of biological sex, EcoHIV treatment upregulated *Syk* and downregulated *Ctnnb1* mRNA expression. Notably, biological sex regulates precisely how EcoHIV treated microglia respond to the suppression of *Trem2* and compensatory hyperproliferative state. Collectively, primary microglia treated with EcoHIV exhibit a highly specialized and sex-dependent transcriptional state. Enhancing our understanding of how HIV-1 alters microglial homeostasis is crucial for the identification of novel therapeutic targets for HAND.

## INTRODUCTION

The human immunodeficiency virus type 1 (HIV-1) pandemic persists, afflicting approximately 40.8 million individuals worldwide [1]. Combination antiretroviral therapy (cART), the standard medical treatment for HIV-1, efficaciously suppresses viral replication in peripheral cells [2]. The blood-brain barrier, however, prevents systemic cART from achieving therapeutic concentrations in the central nervous system (CNS). Consequently, the CNS, which is infiltrated by HIV-1-infected monocytes and CD4+ cells within three to five days of infection [3–4], serves as a major anatomical reservoir for the virus. At the cellular level, microglia, the innate immune cells of the CNS, are the predominant cell type infected by HIV-1 (Clinical: [5–6]; Preclinical: [7–9]). Fundamentally, microglia harbor replication-competent HIV-1 [10–11] supporting an intact cellular viral reservoir.

HIV-1-associated neurocognitive disorders (HAND), which plague approximately 50% of HIV-1 seropositive individuals [12–13], are a direct consequence of the persistent microglial viral reservoir. Clinically, the neurocognitive phenotype of HAND is characterized by profound deficits in attention, memory, and executive function [14]. From a mechanistic perspective, both neuronal [15] and microglial [16–17] dysfunction have been proposed as independent histopathological substrates of HAND. More likely, however, the immediate pathophysiological mechanisms underlying HAND involve a combination of both neuronal injury and defective microglial homeostasis. For example, HIV-1 seropositive individuals with symptomatic HAND exhibit an increased number of microglial nodules [17–18] and a unique microglial transcriptomic signature [18] relative to HIV-1 seronegative persons and/or HIV-1 seropositive individuals without neurological disease. Crucially, microglial dysregulation may directly drive HIV-1-induced synaptodendritic damage [17–19]. Given that microglia are central orchestrators of both HIV-1-induced neurocognitive impairments and neuronal damage, a more comprehensive understanding of how HIV-1 alters microglial homeostasis is crucial for the identification of novel therapeutic targets for HAND.

Preclinical systems, including, but not limited to, single viral protein models (e.g., Transactivator of Transcription (Tat) mice), chimeric HIV (EcoHIV), and simian immunodeficiency virus (SIV) have been instrumental in further characterizing key aspects of HIV-1-induced microglial dyshomeostasis. First, microglia exposed to HIV-1 viral proteins exhibit a sustained proinflammatory state evidenced, in part, by the upregulation of TREM1 [20], as well as genes encoding subunits of the NADPH oxidase family of enzymes [21], those forming a core enzymatic cascade producing prostaglandin E2 [20], and the S100 family of genes [22]. The proinflammatory state exhibited by HIV-1 treated microglia is also driven by the assembly of the NLR family pyrin domain containing 3 (NLRP3) inflammasome [23–25]. Second, the HIV-1 Tat protein promotes microglial survival [20]. Finally, EcoHIV induces profound microglia proliferation characterized by an increased number of microglia colocalized with marker of proliferation Ki-67 (MKI67) relative to controls [26].

Under homeostatic conditions, microglia are long-lived cells (i.e., approximately 4.2 years; [27]) whose density is maintained by an intricate balance of proliferation and apoptosis [28]. Indeed, despite variability in the predicted microglia turnover rate (e.g., Humans: 0.08% to 2% per day [27–28]; Murine: 0.05% to 0.69% per day [28–29], the turnover rate is relatively slow during adulthood. At the molecular level, microglial survival and density maintenance relies on a regulatory trio, including colony stimulating factor 1 receptor (CSF1R), triggering receptor expressed on myeloid cells 2 (TREM2), and DNAX activating protein of 12 kDa (DAP12; for review, [30–31]). Engagement of CSF1R by its ligands (i.e., CSF-1, Interleukin-34 (IL-34)) induces homodimerization of the receptor and activation of downstream signaling pathways. In contrast, activation by phospholipids and lipoproteins via ligand binding, induces a conformational shift in the TREM2/DAP12 complex resulting in the phosphorylation of the immunoreceptor tyrosine-based activation motif (ITAM), thereby activating downstream pathways. Crucially, microglia proliferation is accelerated by neurotoxin exposure (e.g., Incidental Lewy Body Disease: [32]; HIV-1: [26]; Alzheimer’s Disease: [33]). To date, however, the receptors and/or downstream signaling pathways underlying the exacerbated microglial proliferation observed in HIV-1 remain understudied.

To address this fundamental knowledge gap, *in vitro* studies were undertaken. EcoHIV, originally developed by Potash et al. [34], was used in the present study as a biological system to model systemic HIV-1 infection. Crucially, EcoHIV infects microglia [9, 35] by binding to the cationic amino acid transporter (CAT-1) and exacerbates microglia proliferation at the protein level *in vivo* [26]. Herein, primary microglia were treated with EcoHIV and complementary techniques (i.e., quantitative polymerase chain reaction (qPCR) and *in situ* hybridization) were used to evaluate the molecular pathway underlying the exacerbated microglia proliferation induced by EcoHIV at the transcriptomic level. Given the inherent sexual dimorphism present in primary microglia (for review, [36]), the factor of biological sex was integral to the experimental design. Enhancing our understanding of how HIV-1 alters microglial homeostasis is crucial for the identification of novel therapeutic targets for HAND.

## MATERIALS AND METHODS

### Animals

Pregnant female Fischer (F344/NCrl) dams were procured from Charles River Laboratories (Wilmington, MA, USA) during late gestation. To control for independence of observations, the goal was to use primary mixed glia isolated from one rodent pup per sex for each technical replicate (i.e., qPCR or *in situ* hybridization) or gene of interest (i.e., *In Situ* Hybridization: *Pcna*, *Mki67*, *Csf1r*, *Trem2*, *Dap12*).

Rodents had *ad libitum* access to rodent food (2018 Teklad 18% Protein Rodent Diet) and water. Animal husbandry was provided by the University of Kentucky’s Division of Laboratory Animal Resources. The study was conducted in accordance with the 8th Edition of the Institute for Laboratory Animal Research Guide for the Care and Use of Laboratory Animals. Rodents were housed in AAALAC-accredited facilities with environmental conditions targeted at 21°C ± 2°C, 50% ± 10% relative humidity, and a 12-h light:12-h dark cycle. The University of Kentucky Institutional Animal Care and Use Committee approved the project protocol under Federal Assurance (#D16-00217).

### Chimeric HIV (EcoHIV)

#### EcoHIV Virus Construction and Preparation

The EcoHIV-NL4-3-EGFP lentivirus, which was constructed from pNL4-3 and pNCA-WT, were generously provided by Dr. Mary Jane Potash and Dr. David Volsky at the Icahn School of Medicine at Mount Sinai. Additional details regarding the generation of the chimeric EcoHIV proviral plasmid are reported in Potash et al. [34].

#### EcoHIV Plasmid Amplification and Purification

EcoHIV plasmids were transformed into DH5α E. coli using the procedures outlined by Alfar et al. [37] with minor modifications. In brief, 1 μL of the EcoHIV plasmid was added to 50 μL of DH5α E. coli and kept on ice for 30 minutes. The bacteria were heat shocked at 42^◦^C for 90 seconds and once again placed on ice for 2 minutes. The bacteria were spread onto lysogeny broth (LB) agar plates containing ampicillin (100 μg/mL; Thermo Fisher Scientific, Waltham, MA) and incubated overnight at 37°C. Bacteria were amplified by adding a single colony of E. coli to 4 mL ampicillin-containing LB media; media was incubated at 37°C with shaking for six hours at 250 rpm. The 4 mL LB media containing both ampicillin and the E. coli colony were transferred into 1L of ampicillin-containing LB media and shaken overnight (37°C at 250 rpm). The next day, bacteria were harvested by centrifugation for 30 minutes at 4°C. EcoHIV plasmids were subsequently isolated using the HiSpeed Plasmid Maxi Kit (Qiagen, Hilden, Germany). A nanodrop (NanoDrop One/One Microvolume UV-Vis Spectrophotometer, Thermo Fisher Scientific) was used to determine the plasmid concentration.

#### EcoHIV Production

293FT cells (Thermo Fisher Scientific) were cultured in poly-L-lysine coated 75 cm^2^ flasks with medium (i.e., DMEM/F12 Media (Corning, Tewksbury, MA) Supplemented with 10% Fetal Bovine Serum (FBS; Cytiva Life Sciences, Marlborough, MA)). Cells were incubated (37°C, 5% CO^2^/95% Room Air) and passaged at least five times before transfection. For each passage, 293FT cells were incubated with 2-3 mL of 0.25% trypsin-EDTA (GIBCO Life Technologies, Grand Island, NY) until the cells were rounded and beginning to detach; trypsin-EDTA was inactivated with DMEM/F12 media supplemented with 10% FBS. Cells were resuspended in media and replated in 75 cm^2^ flasks.

293FT cells were transfected with EcoHIV using Lipofectamine 3000 Transfection Reagent (Invitrogen by Thermo Fisher Scientific) using the manufacturer’s protocol. Specifically, 22.5 μL of Lipofectamine 3000 Reagent was diluted in 750 μL of Opti-MEM medium (GIBCO). A master mix of DNA was created by mixing 20 μg EcoHIV DNA, 40 μL of P3000 Reagent, and 750 μL of Opti-MEM medium. The DNA master mix and diluted Lipofectamine 3000 Reagent were mixed and incubated at room temperature for 15 minutes. The DNA-lipid complex was added to the 293FT cells and incubated for 48 hours (37°C, 5% CO^2^/95% Room Air). Subsequently, 24 mL of conditional medium containing the packaged lentivirus was centrifuged at 500 g for 10 minutes (4°C); the clarified supernatant was combined with 8 mL Lenti-X Concentrator (Takara Bio, San Jose, CA) and incubated at 4°C. After 48 hours of incubation, the mixture was centrifuged at 1,500 g for 45 minutes (4°C) and the supernatant was removed. The pellet was resuspended in 100 μL of 100 mM phosphate buffered saline (PBS).

### Primary Mixed Glial Cell Cultures

Male and female neonatal (i.e., Postnatal Day 1-3) Fischer F344/NCrl rodent pups were humanely euthanized via decapitation for the isolation of cortical mixed glia. Female Fischer F344/NCrl dams were first offered to other principal investigators, and if immediate placement for terminal procedures was not found, they were humanely euthanized. Specifically, after deeply anesthetizing female dams with inhalant isoflurane (5%), rodents were rapidly decapitated.

After aseptic removal of the brain and meninges from the rodent pup, a piece of the cortex was finely minced with a blade in a small Petri dish containing Hank’s balanced salt solution (HBSS; Corning®) buffered with 10 mM 4-(2-hydroxyethyl)-1-piperasineethanesulfonic acid (HEPES; MilliporeSigma, Burlington, MA). The minced tissue was transferred to a 15 mL centrifuge tube with 2 mL HBSS buffered with 10 mM HEPES and 35 μL 0.25% Trypsin/EDTA and incubated (37°C, 5% CO^2^/95% Room Air) for 20 minutes. The tissue was dissociated using trituration and transferred into a 75 cm^2^ poly-L-lysine coated flask with medium (i.e., DMEM/F12 Media Supplemented with 10% Fetal Bovine Serum (FBS) and 1% Antibiotics (Cytiva)). The culture medium was replaced the day after isolation and subsequently supplemented every three to seven days.

Upon reaching 90% confluency, microglia were purified from primary mixed glial cultures using an orbital shaker. Specifically, flasks were shaken in the incubator for one hour at 220 rpm. After shaking, the culture medium was aspirated and centrifuged for four minutes at 1500 rpm. Microglia were resuspended in culture medium and seeded into either a poly-L-lysine-coated six-well plate or 4-well CulturesSlides (Corning) for qPCR or *in situ* hybridization, respectively. EcoHIV (1 μL per mL of DMEM/F12 Media Supplemented with 10% FBS and 1% Antibiotics) and control (DMEM/F12 Media Supplemented with 10% FBS and 1% Antibiotics) treatment occurred the day after microglia purification.

### Quantitative Polymerase Chain Reaction

#### Cell Preparation

For the primary genes of interest (i.e., *Pcna*, *Mki67*, *Csf1r*, *Trem2*, and *Dap12*), sample sizes were *n*=9 per group (i.e., Male Control, Male EcoHIV, Female Control, Female EcoHIV). Sample sizes for the preliminary analyses of downstream microglial signaling pathways were *n*=5 per group.

Forty-eight hours after control or EcoHIV treatment, one mL of TRIzol® (Invitrogen by Thermo Fisher Scientific) reagent was utilized to lyse microglia directly in the six-well plate. The lysate was homogenized by pipetting up and down several times. Samples were stored at -20°C until further experimentation.

#### RNA Isolation and cDNA Synthesis

The Invitrogen PureLink RNA Mini Kit (Invitrogen by Thermo Fisher Scientific) was utilized to isolate RNA from the homogenized lysate according to the manufacturer’s instructions. A nanodrop (NanoDrop One/One Microvolume UV-Vis Spectrophotometer, Thermo Fisher Scientific) was used to determine the yield of the purified RNA. Complementary DNA was subsequently synthesized from the extracted RNA using Oligo(dT) primer (Promega, Madison, WI, USA) and SuperScript IV Reverse Transcriptase (Invitrogen by Thermo Fisher Scientific).

#### Primer Design

The National Center for Biotechnology Information Primer-Designing Tool and Basic Local Alignment Search Tool database were used to design primer sequences and confirm their specificity, respectively. Custom qPCR primers were manufactured by Integrated DNA Technologies (Coralville, IA, USA). The respective sequences for housekeeping genes and genes are interest are reported in Table 1.

**Table 1.**
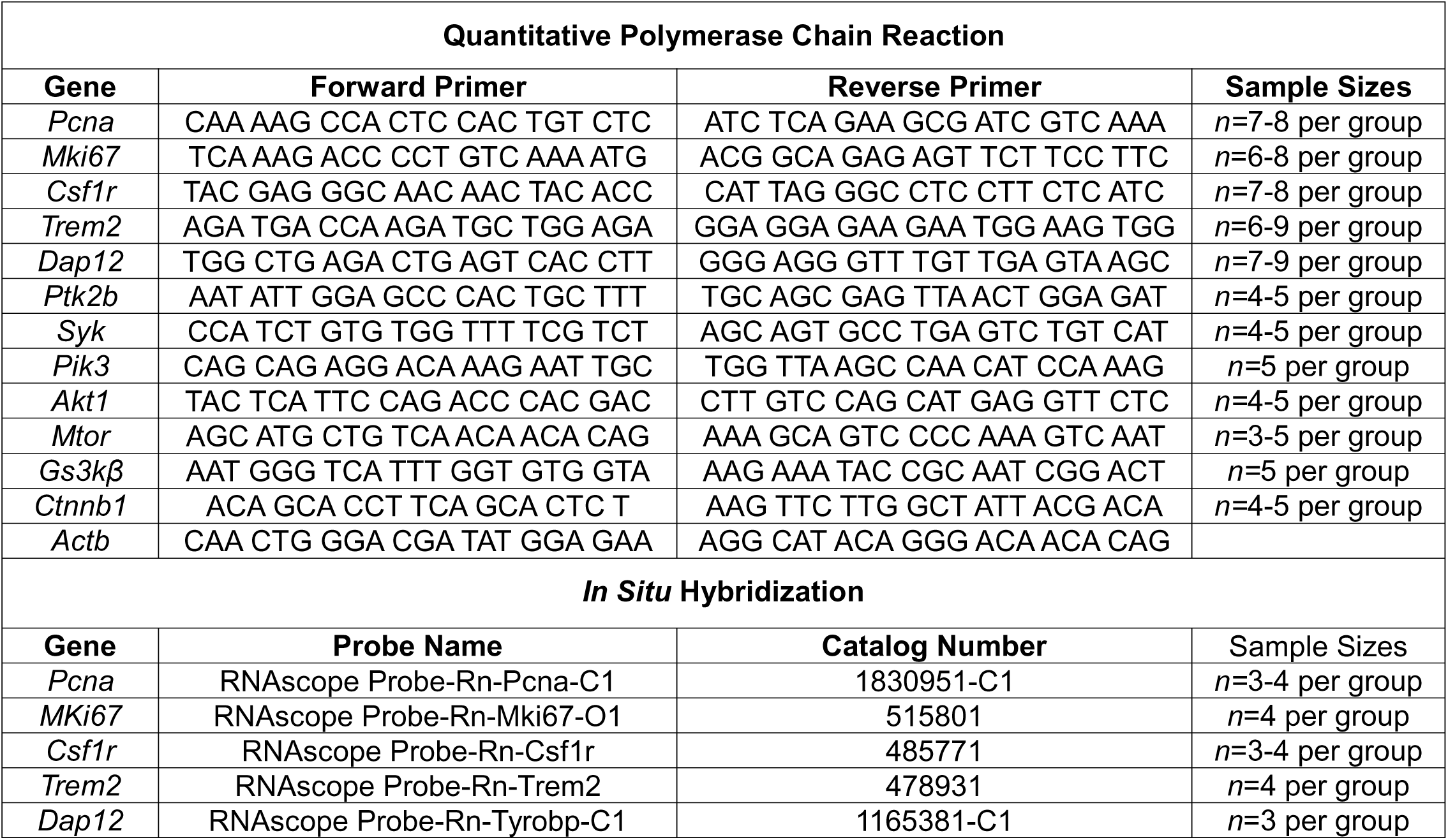
Sequence of primers used in quantitative reverse transcriptase polymerase chain reaction and the highly specific hybridizing probe used for *in situ* hybridization.

#### SYBR-Green qPCR

For each primer evaluated, a primer mixed composed of 20 µm forward primer, 20 µm reverse primer, and nuclease-free water, was prepared in a ratio of 1:1:8. Seven microliter qPCR reactions were assembled in a 384-well plate using 1 µL cDNA, 2 µL of primer mix, and 4 µL of 2x SYBR Green Mix (Applied Biosystems, Waltham, MA, USA). The Bio-Rad CFX384 Real-Time PCR Detection System (Bio-Rad Laboratories, Hercules, CA) was used to perform real-time PCR reactions with the following cycling conditions: 95°C for 10 minutes, 40 cycles of 95°C for 15 seconds followed by 60°C for 1 minute, and a final melt curve of 60°C to 95°C in 0.5°C increments over 10 seconds. The relative gene expression of each gene of interest was analyzed using the 2^-ΔΔCT^ method [38].

Three housekeeping genes, including beta-actin (*Actb*), histone H3.3 (*H3f3b*), and succinate dehydrogenase complex subunit a (*Sdha*), were critically tested. The transcriptomic expression of *H3f3b* and *Sdha* systematically changed across treatment conditions. *Actb*, however, did not systematically change across treatment condition and exhibited strong stability. Thus, target genes were normalized to *Actb*. Gene expression values were normalized to control treated microglia isolated from male pups.

### RNAscope In Situ Hybridization

#### Procedure

Five genes of interest (i.e., *Pcna, Mki67*, *Csf1r*, *Trem2*, *Dap12*) were detected using a highly sensitive and specific *in situ* hybridization technique (Advanced Cell Diagnostics, Inc., Newark, CA, USA). Dependent upon the gene of interest and the factor of biological sex, sample sizes were *n*=3 to *n*=4 per group (i.e., Male Control, Male EcoHIV, Female Control, Female EcoHIV)

Forty-eight hours after control or EcoHIV treatment, culture media was removed and the 4-well CultureSlides were submerged into 1X PBS. Microglia were fixed using 10% neutral buffered formalin (Thermo Fisher Scientific), washed with PBS, and dehydrated using an increasing ethanol (EtOH) gradient (i.e., 50% EtOH for 5 minutes, 70% EtOH for 5 minutes, 100% EtOH for 5 minutes, 100% EtOH for 10 minutes). CultureSlides were stored in 100% EtOH at -20°C until further experimentation.

Upon continuation of the *in situ* hybridization assay, a decreasing EtOH gradient (i.e., 70% EtOH for 2 minutes, 50% EtOH for 2 minutes) and PBS (10 minutes) was used to rehydrate the cells. The ImmEdge Hydrophobic Barrier PAP Pen (Vector Laboratories, Burlingame, CA, USA) was used to create a hydrophobic barrier around each well and allowed to dry. After the slides were rinsed with PBS, each well was incubated with the RNAscope Protease III reagent (Advanced Cell Diagnostics, Inc.) in the HybEZ Humidity Control Tray for 10 minutes. The slides were washed by agitating them in PBS twice.

The RNAscope Multiplex Fluorescent Assay Version 2 (Advanced Cell Diagnostics, Inc.) was subsequently used to conduct *in situ* hybridization. A highly specific hybridizing probe (Table 1) was applied to the wells and CultureSlides were incubated in the HybEZOven at 40°C for 2 hours. After washing twice with RNAscope Wash Buffer, signals were amplified using the “FL v2 AMP 1”, “FL v2 AMP2”, and FL v2 AMP3” reagents provided in the RNAscope Multiplex Fluorescence Assay Version 2 Kit. Wells were covered with the reagents and CultureSlides were incubated in the HybEZ Oven for either 30 minutes (“FL v2 AMP1”, “FL v2 AMP2”) or 15 minutes (“FL v2 AMP3”) at 40°C. Subsequently, the fluorescent signal was developed by sequentially incubating the CultureSlides with “FL v2 HRP-C1" (Advanced Cell Diagnostics, Inc.; 15 minutes), TSA Vivid Fluorophore 570 (Advanced Cell Diagnostics, Inc.) diluted with TSA buffer (Advanced Cell Diagnostics, Inc.; 1:1500; 30 minutes), and “FL v2 HRP Blocker” (Advanced Cell Diagnostics, Inc.; 15 minutes) in the HybEZOven at 40°C. Finally, the wells were counterstained with DAPI for 30 seconds prior to mounting with EverBrite mounting medium (Biotium, Fremont, CA) and coverslipping.

#### Imaging and mRNA Quantification

DAPI counterstaing was utilized to identify high confluency areas for imaging with the goal of imaging at least 10 cells per well. Images were captured at 60x using an oil-immersion objective on a Nikon AXR inverted confocal microscope system controlled by Version 6.10 of Nikon’s NIS-Elements C software. Fluorophore excitation of the genes of interest (i.e., *Pcna, Mki67*, *Csf1r*, *Trem2*, *Dap12)* and DAPI staining were accomplished using a laser wavelengths of 561 nm and 405 nm, respectively.

Individual cells were cropped from each image to 150 x 150 pixels, blinded to prevent experimental bias, and analyzed using ImageJ software [39]. Particle analysis was utilized to count the number of positive pixels within each individual cell. Each cell was assigned a score ranging from 1 to 8 based on the number of pixels; higher scores reflect a greater number of positive pixels within the cell (Supplementary Figures 1-2).

### Statistical Analysis

Analysis of variance (ANOVA) and linear regression techniques were utilized for all statistical analyses. Figures were created in GraphPad Prism 10. Probability values (i.e., *p* values) and generalized eta squared (η_G_^2^; [40]) were used to evaluate statistical and practical significance, respectively. Statistical significance was established at an α criterion of *p* ≤ 0.05. Practical significance was established at a criterion of η_G_^2^ ≥ 0.14, which is consistent Cohen’s [41] definition of a large effect.

qPCR data was analyzed using a univariate ANOVA (SPSS Statistics 31, IBM Corporation, Somer, NY) with treatment (i.e., Control vs. EcoHIV) and biological sex (i.e., Male vs. Female) serving as between-subjects factors. The delta Ct value was used as the dependent variable of interest to preclude violation of standard statistical assumptions (e.g., normality, homoscedasticity) often violated by the analysis of percentage change data. For *Trem2*, the fold change, rather than the delta Ct value, was used for analysis to mitigate the violation of the assumption of normality. Outliers were detected by calculating a z-score, whereby a data point greater than two standard deviations of the mean were considered outliers. Sample sizes after the removal of outliers are indicated in Table 1. *In situ* hybridization data were analyzed using non-linear regression (GraphPad Prism 10 Software, Inc.).

The utility of microglial proliferation markers (i.e., *Pcna, Mki67*) and/or microglial receptors and adapter proteins (i.e., *Csf1r, Trem2,* and *Dap12*) to correctly classify microglia based on their treatment was evaluated using an exploratory discriminant function analysis (DFA; SPSS Statistics 31). If at least one discriminating variable was previously identified as an outlier, the sample was not included in the DFA. As such, the DFA included the following sample sizes (Microglia Proliferation Markers: Control, *n*=12, EcoHIV, *n*=12; Microglial Receptors and Adapter Proteins: Control, *n*=15, EcoHIV, *n*=16). A discriminant score was calculated for each sample by the statistical software (i.e., SPSS Statistics 31) and utilized for subsequent regression analyses (GraphPad Prism 10). Two influential points, which had the lowest *X* values (i.e., Discriminant Score: Microglia Proliferation) were removed from the linear regression.

## RESULTS

### EcoHIV treatment exacerbates microglial proliferation evidenced by increased transcriptomic expression of *Pcna* and *Mki67*

The transcriptomic expression of *Pcna* and *Mki67*, widely used markers of cellular proliferation, was investigated using two complementary approaches, including qPCR and *in situ* hybridization.

*Pcna.* mRNA expression, measured using qPCR, of *Pcna* was significantly increased in in primary microglia treated with EcoHIV relative to control treated cells (Figure 1A; Main Effect of Treatment [*F*(1,25)=10.8, *p*≤0.003, η_G_^2^=0.283]).

**Figure 1.**
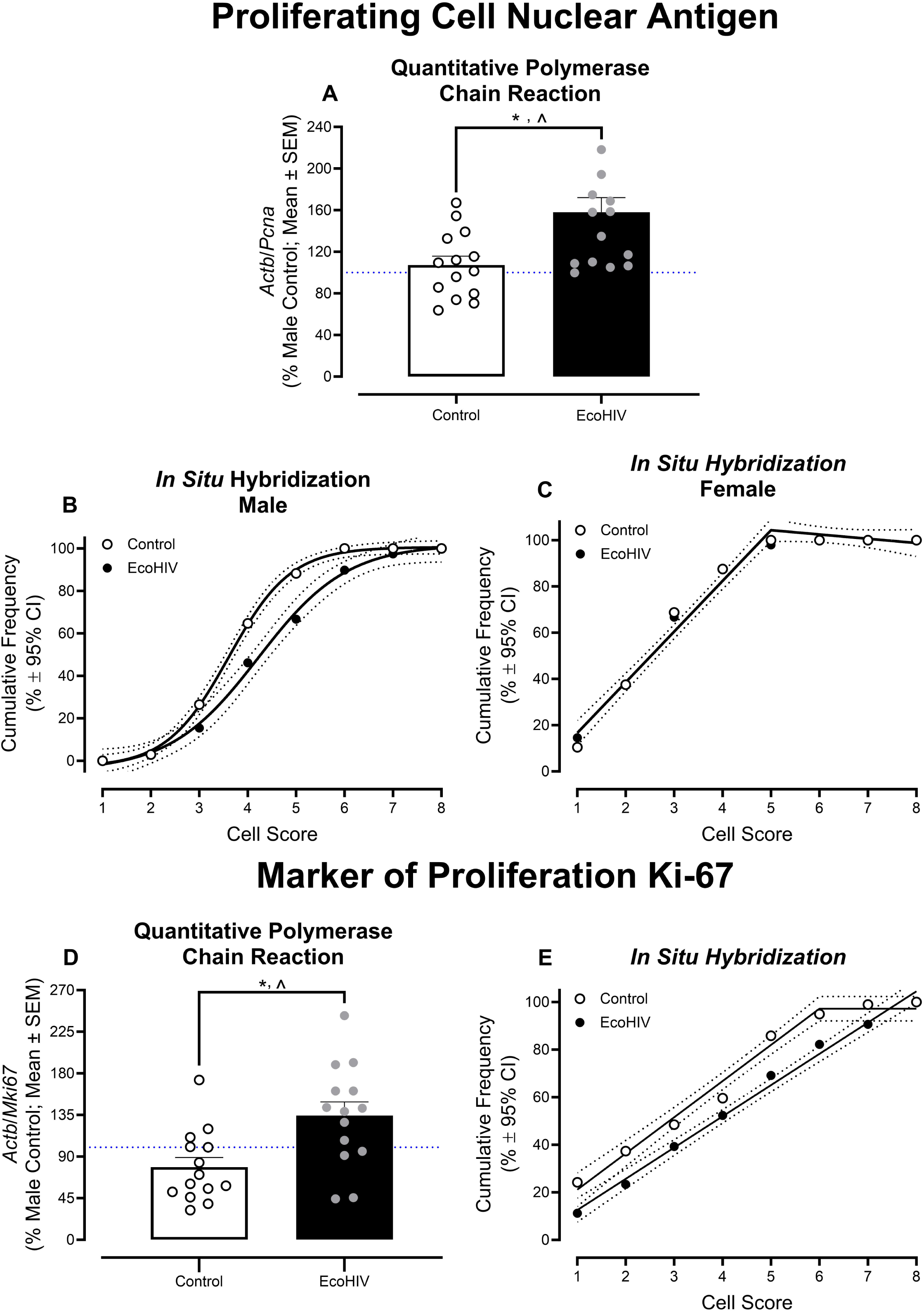
Chimeric HIV (EcoHIV) treatment induces profound transcriptomic increases in markers of microglia proliferation. (**A-C**) In quantitative polymerase chain reaction (qPCR) assessments, the transcriptomic expression of proliferating cell nuclear antigen (*Pcna*) was significantly upregulated by treatment with EcoHIV. *In situ* hybridization measures revealed a significant increase in the frequency of cells in higher-order categories in EcoHIV treated microglia isolated from male pups relative to their control counterparts. EcoHIV treatment, however, failed to significantly alter the cumulative *Pcna* cell score distribution in microglia isolated from female rats. (**D-E**) Independent of biological sex, qPCR analyses revealed a significant EcoHIV treatment induced increase in marker of proliferation Ki-67 (*Mki67*) mRNA expression; observations which were confirmed using *in situ* hybridization. The blue dotted line represents 100% mRNA expression for control male animals. \**p*<0.05 ^ η_G_^2^ > 0.14

*In situ* hybridization (Supplementary Figure 1A) in concert with linear regression analyses revealed treatment- and sex-dependent alterations in the transcriptomic expression of *Pcna*. Specifically, EcoHIV-treated microglia isolated from male rodents exhibited an increased frequency of cells in higher-order categories relative to their control counterparts (Figure 1B). The cumulative cell score distribution for both control and EcoHIV treated microglia was well-described by a sigmoidal dose-response curve with a variable slope (*R*^2^≥0.99) albeit statistically significant differences in the best-fit function were observed [*F*(4,8)=20.9, *p*≤0.001]. In contrast, treatment failed to significantly alter the cumulative *Pcna* cell score distribution in microglia isolated from female rats (Figure 1C; Global Best Fit Function: One-Phase Association, *R*^2^≥0.98).

#### Mki67

Independent of biological sex, transcriptomic expression of *Mki67,* indexed using qPCR, was significantly increased in primary microglia treated with EcoHIV relative to control treated cells (Figure 1D; Main Effect of Treatment [*F*(1,24)=8.8, *p*≤0.007, η_G_^2^=0.224]). Quantification of *in situ* hybridization (Supplementary Figure 1B) in concert with linear regression analyses confirmed these observations, whereby EcoHIV-treated microglia displayed an increased frequency of cells in higher-order categories relative to their control counterparts (Figure 1E). Indeed, the cumulative cell score distribution for control and EcoHIV-treated microglia was well-described by a segmental linear regression (*R*^2^≥0.98) and first-order polynomial (*R*^2^≥0.99), respectively. Notably, the factor of biological sex influenced the magnitude, but not direction, of *Mki67* genotypic dependent alterations observed using *in situ* hybridization techniques (Data Not Shown).

### Transcriptomic expression of microglial receptors and adapter proteins is dysregulated by treatment with EcoHIV

Two crucial microglia receptors (i.e., *Csf1r, Trem2* (Figure 2)) and one transmembrane signaling adapter protein (i.e., *Dap12* (Figure 3)) form an interconnected signaling triad that synergistically regulates microglial proliferation in the CNS. As such, *Csf1r*, *Trem2*, and *Dap12* were subsequently investigated to evaluate the mechanism underlying the exacerbated microglia proliferation induced by treatment with EcoHIV.

**Figure 2.**
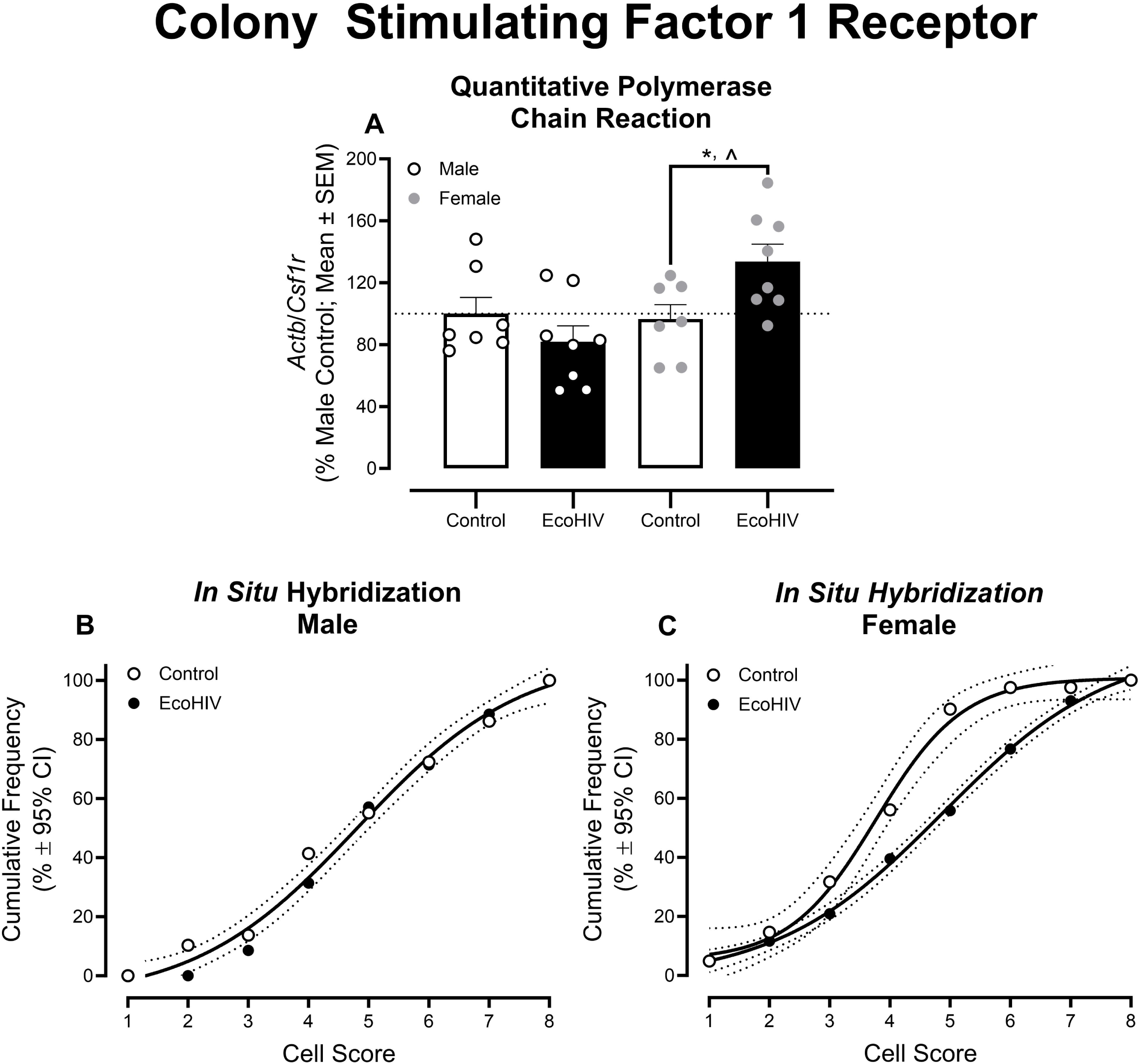
The mRNA expression of colony stimulating factor 1 receptor (*Csf1r*) was sex-dependently upregulated by chimeric HIV (EcoHIV) treatment. (**A**-**C**) Quantitative polymerase chain reaction (qPCR) techniques and *in situ* hybridization revealed an upregulation of *Csf1r* transcriptomic expression in EcoHIV treated microglia isolated from female rodents. The blue dotted line represents 100% mRNA expression for control male animals. \**p*<0.05 ^ η_G_^2^ > 0.14

**Figure 3.**
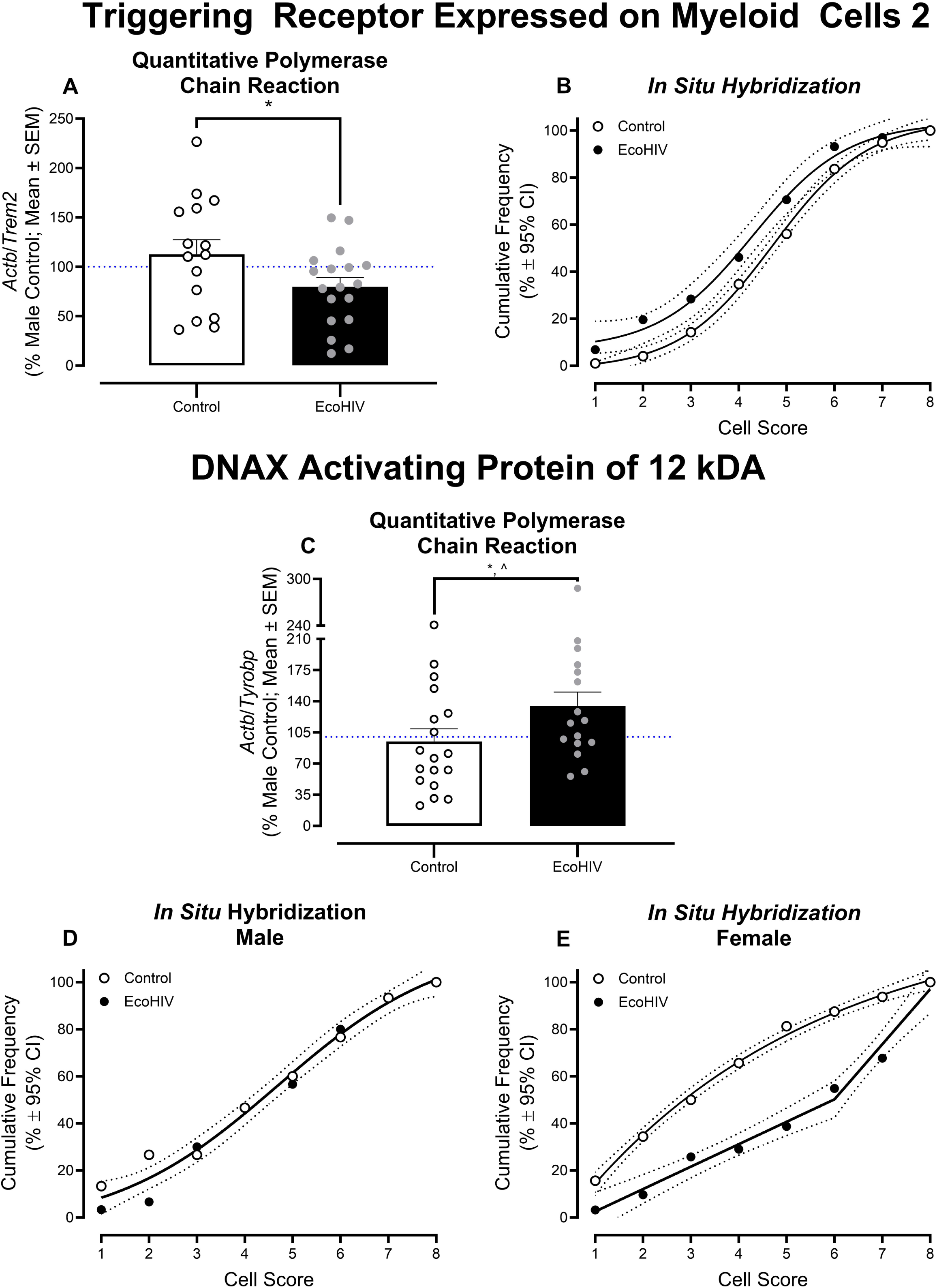
The transcriptomic expression of triggering receptor expressed on myeloid cells 2 (*Trem2*) and DNAX activating protein of 12 kDA (*Dap12*) was altered by treatment with chimeric HIV (EcoHIV) (**A-B**) Independent of biological sex, *Trem2* was significantly downregulated by EcoHIV treatment evidenced by quantitative polymerase chain reaction (qPCR) techniques and *in situ* hybridization. (**C-E**) In sharp contrast, qPCR analyses revealed a significant increase in *Dap12* transcriptomic expression. Nevertheless, *in situ* hybridization measures revealed sex-dependent alterations. Indeed, a significant increase in the frequency of cells in higher-order categories was observed in EcoHIV treated microglia isolated from female pups relative to their control counterparts. The blue dotted line represents 100% mRNA expression for control male animals. \**p*<0.05 ^ η_G_^2^> 0.14

#### Csf1r

qPCR approaches revealed a treatment- and sex-dependent alteration in the transcriptomic expression of *Csf1r* in primary microglia (Figure 2A; Treatment x Sex Interaction [*F*(1,26)=7.1, *p*≤0.013, η_G_^2^=0.183]). Complementary analyses were conducted by sex to elucidate the locus of the interaction. Indeed, EcoHIV treated microglia isolated from female pups exhibited a statistically significant increase in *Csf1r* mRNA expression relative to their control counterparts (Main Effect of Treatment: [*F*(1,13)=6.3, *p*≤0.026, η_G_^2^=0.326]). EcoHIV treatment failed to significantly alter *Csf1r* transcriptomic expression in microglia isolated from male rats (*p*>0.05 and η_G_^2^<0.14).

*In situ* hybridization (Supplementary Figure 2A) in concert with linear regression analyses (Figure 2B-C) confirmed these observations. A sigmoidal dose-response curve with a variable slope provided the best-fit function (*R*^2^≥0.99) for the cumulative cell score distribution for both EcoHIV and control treated microglia isolated from female rodents; albeit statistically significant differences in the fit of the function were observed [*F*(4,8)=29.3, *p*≤0.001]. Treatment, however, had no statistically significant effect on *Csf1r* cell score in microglia isolated from male rats evidenced by a global best fit function (i.e., Sigmoidal Dose-Response Curve with a Variable Slope, *R*^2^≥0.99).

#### Trem2

Independent of biological sex, mRNA expression of *Trem2,* indexed using qPCR, was significantly decreased in EcoHIV, relative to control, treated primary microglia (Figure 3A; Main Effect of Treatment [*F*(1,29)=4.7, *p*≤0.039, η_G_^2^=0.125]). Quantification of *in situ* hybridization (Supplementary Figure 2B) confirmed these observations, whereby a prominent leftward population shift,supporting an increased frequency of cells in lower-order categories, was observed in EcoHIV treated, relative to control treated, microglia (Figure 3B). The cumulative cell score distribution was well-described by a sigmoidal dose-response curve with a variable slope (*R*^2^≥0.99) albeit statistically significant treatment-induced differences in the best-fit function were observed [*F*(4,8)=12.2, *p*≤0.002].

#### Dap12

Similarly, qPCR results support an EcoHIV induced increase in the transcriptomic expression of *Dap12* independent of biological sex (Figure 3C; Main Effect of Treatment [*F*(1,30)=5.3, *p*≤0.029, η_G_^2^=0.146]).

*In situ* hybridization (Supplementary Figure 2C) in concert with linear regression analyses, however, revealed treatment- and sex-dependent alterations in the transcriptomic expression of *Dap12* (Figure 3D-E). The cumulative cell score distribution of primary microglia isolated from female rodents was well-described by a one-phase association (*R*^2^≥0.99) and segmental linear regression (*R*^2^≥0.98) following control or EcoHIV treatment, respectively. As such, EcoHIV treatment induced an increased frequency of microglia in higher-order categories relative to control treatment. Treatment, however, had no statistically significant effect on the cumulative *Dap12* cell score distribution in microglia isolated from male rats (Global Best Fit Function: Sigmoidal Dose-Response Curve with a Variable Slope, *R*^2^≥0.99).

### Dysregulation of the Trem2/Dap12 complex is strongly associated with the exacerbated microglia proliferation induced by EcoHIV

An exploratory discriminant function analysis was utilized to determine which markers of cellular proliferation (i.e., *Pcna*, *Mki67*) were best able to identify microglia treatment. A stepwise discriminant function analysis selected two variables (*Pcna* delta Ct and *Mki67* delta Ct) that maximally separated EcoHIV and control microglia (Figure 4A; Canonical Correlation of 0.629). Microglia cells were classified (jackknifed) with 79.2% accuracy (*F* approximation of Wilks’ λ of 0.605, [*F*(2,22)=6.9, *p*≤0.005]).

**Figure 4.**
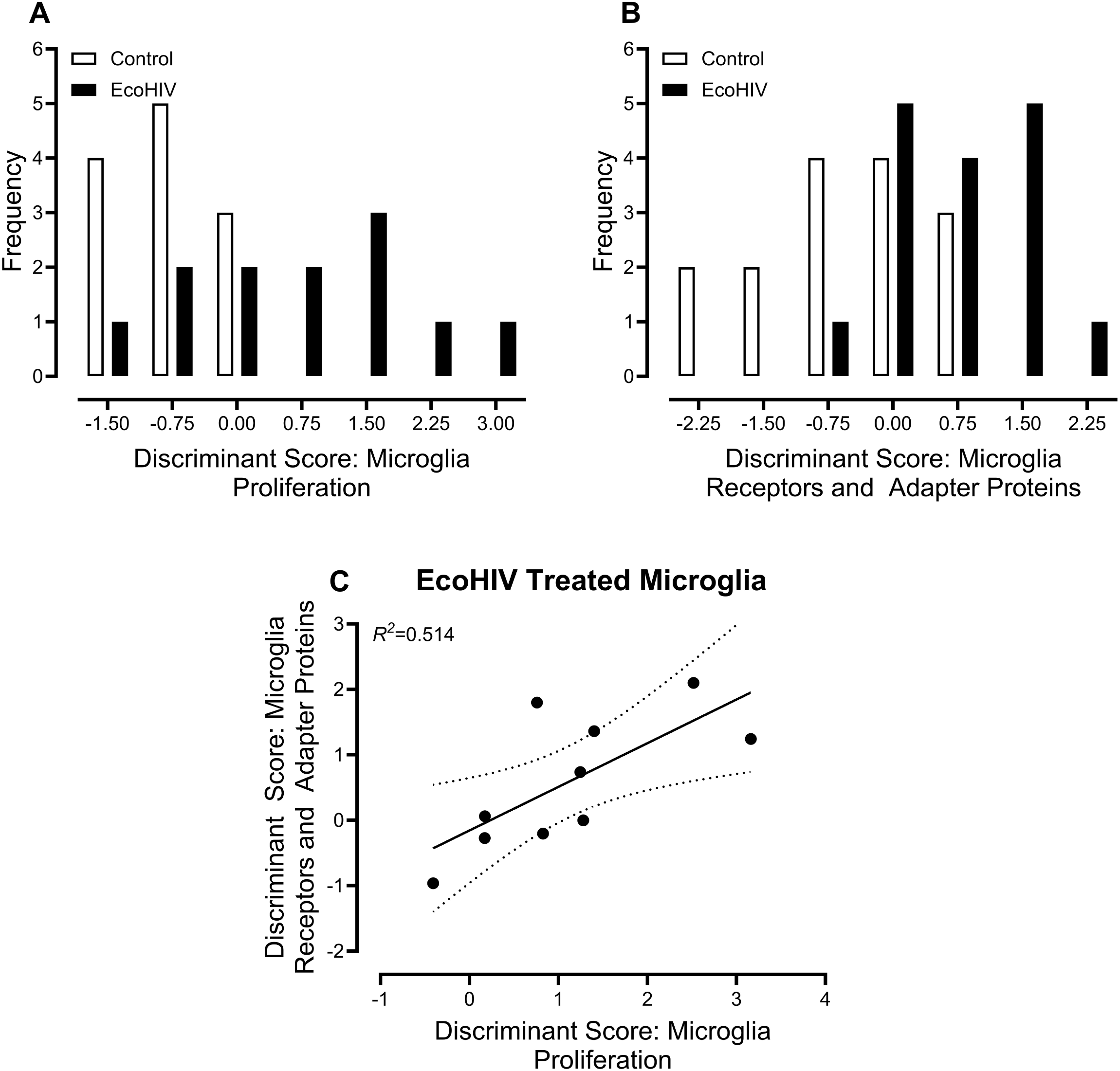
EcoHIV induced dysregulation of triggering receptor expressed on myeloid cells 2 (*Trem2*) and DNAX activating protein of 12 kDA (*Dap12*) are strongly associated with microglia proliferation. (**A**) A discriminant function analysis (DFA) was utilized to determine whether measures of microglia proliferation could classify animals based on treatment. Microglia cells were classified (jackknifed) based on treatment with 79.2% accuracy using proliferating cell nuclear antigen (*Pcna*) and marker of proliferation Ki-67 (*Mki67*) delta Ct values. (**B**) Subsequently a DFA was utilized to determine whether measures of microglia receptors (i.e., colony stimulating factor 1 receptor (*Csf1r*), *Trem2*) and/or adapter proteins (i.e., *Dap12*) could classify microglial cells based on treatment. Microglia cells were classified (jackknifed) based on treatment with 71.0% accuracy using *Trem2* and *Dap12* delta Ct values. (**C**) Finally, a linear regression analysis was conducted to evaluate if the dysregulation of microglial receptors and adapter proteins is associated with microglial proliferation in EcoHIV treated microglia; both microglial receptors and adapter proteins, as well as microglial proliferation, were indexed using discriminant scores. In EcoHIV-treated microglia, 51.4% of the variance in microglial proliferation was accounted for by microglial receptors and adapter proteins.

A second exploratory discriminant function analysis was conducted to evaluate which microglial receptors and adapter proteins (i.e., *Csf1r*, *Trem2*, *Dap12*) were best able to identify microglia treatment. A stepwise discriminant function analysis selected two variables (i.e., *Trem2* delta Ct and *Dap12* delta Ct) that maximally separated EcoHIV and control microglia (Figure 4B; Canonical Correlation of 0.560). Microglia cells were classified (jackknifed) with 71.0% accuracy (*F* approximation of Wilks’ λ of 0.687, [*F*(2,23)=5.02, *p*≤0.016]).

A discriminant score was calculated for each sample by the statistical software (i.e., SPSS Statistics 31) and utilized for subsequent regression analyses. Indeed, a linear regression analysis was conducted to evaluate if the dysregulation of microglial receptors and adapter proteins is associated with microglial proliferation in EcoHIV treated microglia; both microglial receptors and adapter proteins, as well as microglial proliferation, were indexed using discriminant scores. In EcoHIV treated microglia, 51.4% of the variance in microglial proliferation was accounted for by microglial receptors and adapter proteins (Figure 4C). Exploratory analyses, therefore, support a strong association between the dysregulated Trem2/Dap12 complex and microglial proliferation in EcoHIV.

### A preliminary investigation of downstream microglial signaling pathways supports the induction of a unique transcriptomic profile by EcoHIV

Given the profound association between the dysregulated *Trem2*/*Dap12* complex and microglial proliferation in EcoHIV, a preliminary investigation of the Trem2-mediated microglial survival and immunoregulation pathway was conducted. Specifically, a linear hierarchical cascade of genes was explored to map a sequential signal flow from the microglial cell membrane to the nucleus.

#### Membrane-Associated Upstream Kinases

TREM2 and a DAP12 homodimer are constitutively bound via an electrostatic salt bridge. Extracellular ligand binding induces a conformational shift in the TREM2/DAP12 complex resulting in the phosphorylation of the immunoreceptor tyrosine-based activation motif (ITAM). Phosphorylation of the ITAM recruits spleen tyrosine kinase (SYK), which, upon activation, triggers downstream mediators (e.g., Protein Tyrosine Kinase 2 Beta (PTK2B)).

Independent of biological sex, *in vitro* treatment of microglia with EcoHIV significantly increases *Syk* transcriptomic expression (Figure 5A; Main Effect of Treatment: [*F*(1,14)=5.1, *p*≤0.04, η_G_^2^=0.195]). mRNA expression of *Ptk2b*, however, was altered in a treatment and sex dependent manner (Figure 5B; Treatment x Sex Interaction: [*F*(1,14)=7.5, *p*≤0.016, η_G_^2^=0.304]). Complementary analyses were conducted by sex to identify the locus of the interaction. Indeed, a practically (i.e., η_G_^2^=0.343 – 0.378), but not statistically (i.e., *p*≤0.098), significant increase or decrease in *Ptk2b* was observed in EcoHIV treated microglia isolated from male and female rat pups, respectively (Main Effect of Treatment (Male): [*F*(1,7)=4.3, *p*≤0.078, η_G_^2^=0.378; Main Effect of Treatment (Female): [*F*(1,7)=3.7, *p*≤0.098, η_G_^2^=0.343]).

**Figure 5.**
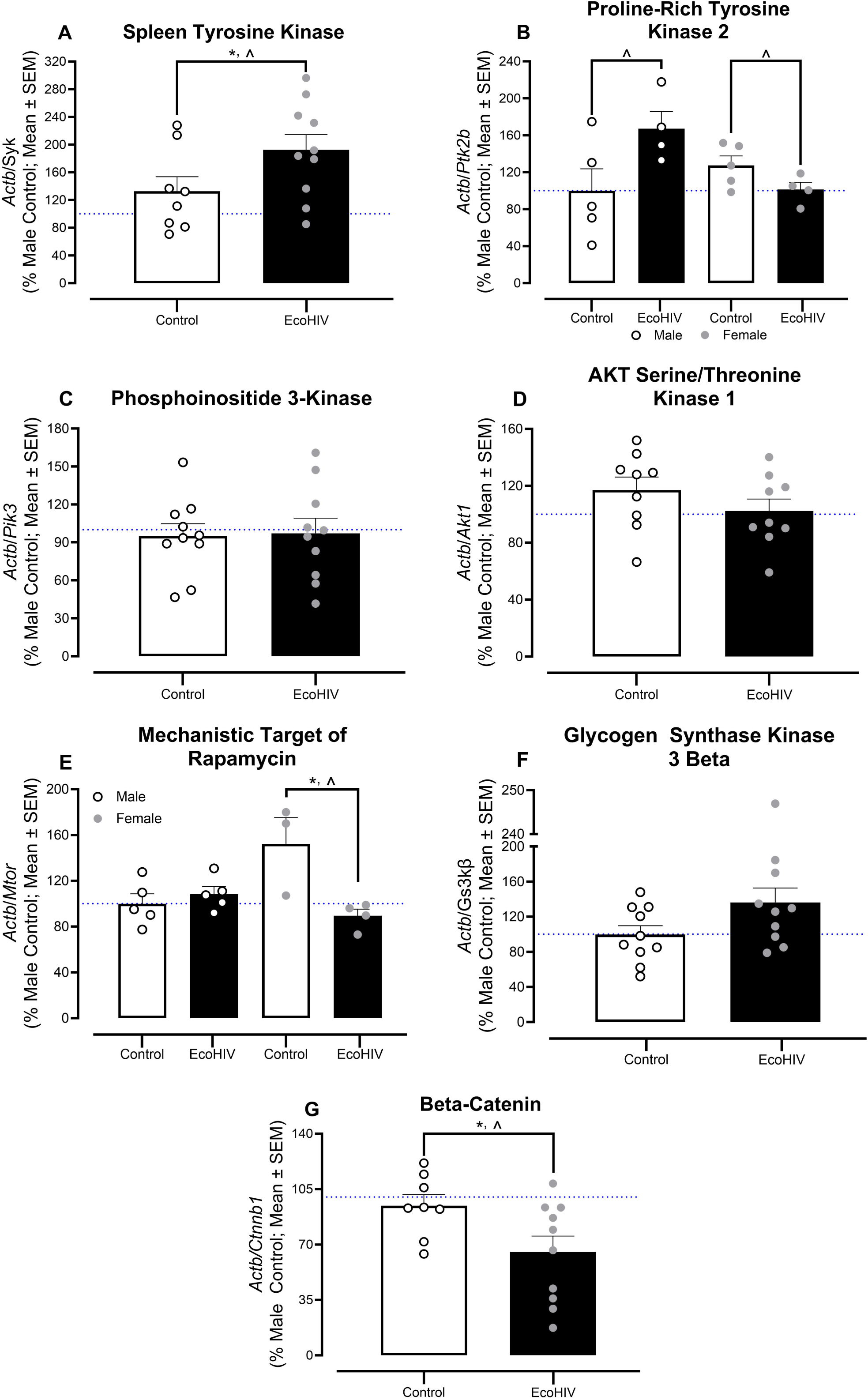
A preliminary investigation of downstream microglial signaling pathways supports the induction of a unique transcriptomic profile by chimeric HIV (EcoHIV). (**A-G**) Independent of biological sex, EcoHIV treatment upregulated the expression of spleen tyrosine kinase (*Syk*) and suppressed the expression of beta-catenin (*Ctnnb1*). Furthermore, in microglia isolated from male rodents, EcoHIV treatment increased the mRNA expression of proline-rich tyrosine kinase 2 (*Ptk2b*). In microglia isolated from female pups, EcoHIV treatment decreased the transcriptomic expression of *Ptk2b* and mechanistic target of rapamycin (*Mtor*). The blue dotted line represents 100% mRNA expression for control male animals. \**p*<0.05 ^ η_G_^2^ > 0.14

#### Phosphoinositide 3-Kinase/AKT Serine/Threonine Kinase 1 Survival Axis

The membrane-associated upstream kinases (e.g., SYK, PTK2B) recruit and activate Class I phosphoinositide 3-kinases (PI3K). The activation of PI3K shifts the signal from a protein-kinase cascade to a lipid-messenger cascade, whereby lipid messengers (e.g., phosphatidylinositol (3,4,5)-trisphosphate (PIP_3_)) activate AKT serine/threonine kinase 1 (AKT) to trigger microglial growth and proliferation.

At the transcriptomic level, EcoHIV treatment failed to alter the *Pi3k*/*Akt* survival axis. Specifically, the mRNA expression of neither *Pi3k* (Figure 5C) nor *Akt* (Figure 5D) was significantly altered by EcoHIV treatment (i.e., *p*>0.05, η_G_^2^<0.14).

Nevertheless, the activation of AKT by PI3K results in the phosphorylation of two distinct downstream targets, including the mechanistic target of rapamycin (mTOR) and glycogen synthase kinase-3 beta (GSK3β); GSK3β, in turn, phosphorylates β-catenin.

Crucially, the transcriptomic expression of *mTOR* was altered in a treatment and sex-dependent manner (Figure 5E; Treatment x Sex Interaction: [*F*(1,13)=11.1, *p*≤0.005, η_G_^2^=0.357), whereby *mTOR* mRNA expression was significantly upregulated by EcoHIV in microglia isolated from female (Main Effect of Treatment: (*F*(1,5)=10.4, *p*≤0.023, η_G_^2^=0.675]), but not male (*p*>0.05, η_G_^2^<0.14) pups. Furthermore, although EcoHIV failed to significantly alter the mRNA expression of GSK3β (Figure 5F; *p*>0.05, η_G_^2^<0.14), treatment significantly downregulated mRNA expression of *Ctnnb1* (Figure 5G; Main Effect of Treatment: [*F*(1,14)=5.3, *p*≤0.023, η_G_^2^=0.231]).

## DISCUSSION

Purified microglia treated with EcoHIV exhibit a highly specialized and sex-dependent transcriptional state. From a functional perspective, treatment with EcoHIV induces profound microglia hyperproliferation evidenced by the upregulation of both *Pcna* and *Mki67* mRNA in qPCR analyses; the exacerbation of microglial proliferation at the mRNA level confirms and extends *in vivo* findings in EcoHIV rats at the protein level [26]. Independent of biological sex, EcoHIV induces a downregulation of *Trem2* and *Ctnnb1*, as well as an upregulation of *Dap12* and *Syk*. Furthermore, treatment- and sex-dependent transcriptional alterations were observed in *Csf1r, Ptk2b,* and *Mtor*. Minor discrepancies between qPCR and *in situ* hybridization, which were observed for *Pcna* and *Dap12*, may afford critical insight into precisely how transcriptomic expression is altered by EcoHIV. Crucially, independent of biological sex, alterations in the transcriptomic expression of *Trem2* and *Dap12* were strongly associated with the exacerbated microglia proliferation induced by EcoHIV. Enhancing our understanding of how HIV-1 alters microglial homeostasis is crucial for the identification of novel therapeutic targets for HAND.

Historically, microglia have been classified using one of two binary frameworks. First, the seminal work of Streit, Graeber, and Kreutzberg [42] categorized microglia as resting or activated based on their morphological characteristics. Indeed, resting microglia were characterized by small cell bodies with highly branched, long processes; the morphological characteristics of activated microglia, in contrast, include a round, amoeboid cell body and retracted, thick processes. Modern technological advances, including non-invasive two-photon *in vivo* imaging of microglia [43–44], however, demonstrated that microglia are continually surveying their microenvironment under homeostatic conditions [43–44]. As such, microglia are never “resting” [45]. Furthermore, traumatic brain injury does not induce “activation” of microglia, per se, but rather alters the nature of their surveillance (e.g., decreased motility of microglial processes; [43–44]). Second, a functional binary model was proposed [46], whereby microglia were categorized as either pro-(M1) or anti-inflammatory (M2) based on gene expression. Single-cell RNA sequencing, however, again revolutionized neuroimmunology by revealing the co-expression of traditional “M1” and “M2” genes within a single cell [47]. As such, the relevance of exploring highly specialized transcriptional states induced by EcoHIV infection cannot be understated.

Indeed, herein, complementary widely used markers of cellular proliferation (e.g., PCNA: [48–49]; MKI67: [50]) were used to investigate how primary microglia respond to treatment with EcoHIV. PCNA, a nuclear protein fundamentally involved in deoxyribonucleic acid (DNA) replication, is highly expressed during the transition from the G1 to S phase of the cell cycle [51]; its expression, however, persists in non-dividing cells due to its longer half-life (i.e., ∼20 Hours; [52]) and function (for review, [53]). In sharp contrast, Ki-67 acts as a physical surfactant to keep mitotic chromosomes separated during cell division [54] and is expressed exclusively in proliferating cells [55–56]. Independent of biological sex, qPCR analyses revealed the upregulation of both *Pcna* and *Mki67* in EcoHIV treated microglia, thereby providing crucial cross-validation of observations across two distinct cell cycle genes. It is notable, however, that *in situ* hybridization revealed sex- and treatment-dependent alterations in the transcriptomic expression of *Pcna*.

On the surface, differences in the conclusions drawn from qPCR and *in situ* hybridization data, which both evaluate mRNA expression, appear contradictory. The techniques, however, are complementary, whereby qPCR provides high-throughput validation of bulk expression levels and *in situ* hybridization affords an opportunity to identify cell-to-cell heterogeneity. Discrepancies between qPCR and *in situ* hybridization analyses, therefore, may actually afford critical insight into precisely how transcriptomic expression is altered by EcoHIV. Indeed, purified microglia isolated from male rodents and treated with EcoHIV may be responding with a robust, highly concentrated transcriptional response that is identified by both qPCR and *in situ* hybridization. In contrast, EcoHIV treated primary microglia isolated from female pups may be exhibiting an upregulation of *Pcna* mRNA at a low level per cell; a transcriptional response that is recognized by qPCR, but not *in situ* hybridization, analyses. Nevertheless, *in vitro* findings from the present study are consistent with previous reports in HIV-1 seropositive individuals with and without encephalitis [57–58], rhesus macaques infected with simian immunodeficiency virus (SIV; [58]) and EcoHIV infected rats *in vivo* [26] supporting pronounced EcoHIV induced microglia hyperproliferation.

Independent of biological sex, qPCR approaches support a stoichiometric imbalance in *Trem2* and *Dap12* by EcoHIV treatment. Specifically, the *in vitro* purified microglia transcriptomic profile in EcoHIV-treated cells is characterized by the downregulation and upregulation of *Trem2* and *Dap12*, respectively; the downregulation of *Trem2* is consistent with prior preclinical [59] and clinical [60] studies. The fundamental import of EcoHIV induced *Trem2* and *Dap12* mRNA alterations cannot be understated. First, herein, a strong association between EcoHIV induced *Trem2* and *Dap12* transcriptomic dysregulation and microglia hyperproliferation was observed. Second, TREM2 is a cell surface receptor that binds to specific anionic ligands (e.g., Lipids, Apolipoproteins [61], Amyloid-Beta [62]). The consequences of *Trem2* downregulation, therefore, may extend beyond microglia hyperproliferation, as the suppression of TREM2 expression accelerates the accumulation of amyloid beta [62–63]. Indeed, with regards to HIV-1, clinical and preclinical studies have shown that HIV induces an abnormal accumulation of intraneuronal amyloid beta [64–66], as well as strong, sex-dependent associations between intraneuronal amyloid beta and synaptic dysfunction [66]. In addition, in HIV-1 seropositive individuals, cerebral spinal fluid concentrations of soluble TREM2 was highly associated (*r*=0.62) with a measure of neuronal injury (i.e., CSF Neurofilament Light Protein; [67]). TREM2, therefore, may serve as a promising therapeutic target for HAND.

EcoHIV treatment also upregulates *Syk* and downregulates *Ctnnb1* (i.e., the gene encoding β-catenin) in purified microglia isolated from male and female rodents. Under homeostatic conditions, phosphorylation of the DAP12 ITAM recruits SYK to activate the PI3K/AKT pathway and stabilize β-catenin; canonical Wnt/β-catenin signaling is a key regulatory pathway promoting microglia survival and proliferation. At first glance, the simultaneous upregulation of *Pcna* and *Mki67*, in combination with *Trem2* and *Ctnnb1* deficiency appears paradoxical. However, under pathological conditions, as is observed in Alzheimer’s disease, microglia compensate for the absence of TREM2 or DAP12 by dysregulating downstream signaling transduction [68]. Additionally, the concurrent downregulation of *Trem2*, in combination with the upregulation of *Dap12* and *Syk,* may suggest that orphan DAP12 proteins couple with other microglial surface receptors (e.g., TREM1 [69], SIRPβ1) to induce a hyperproliferative and inflammatory state.

Crucially, the factor of biological sex regulates precisely how EcoHIV treated microglia respond to the suppression of *Trem2* and compensatory hyperproliferative state. Indeed, EcoHIV treated microglia isolated from male rodents exhibited an increase in *Pyk2* transcriptomic expression. Overexpression of *Pyk2*, in combination with the upregulation of *Dap12* and *Syk* may, at least in part, underlie NLRP3 inflammasome activation [70]; NLRP3 inflammasome activation is a well-recognized consequence of HIV-1 infection [23–25]. In sharp contrast, EcoHIV treated microglia isolated from female rodents exhibited an upregulation of *Csf1r* and a suppression of *Pyk2* and *Mtor*. First, the increased mRNA expression of *Csf1r*, which confirms and extends prior research by considering the factor of biological sex [71–72], may reflect another way that EcoHIV treated microglia isolated from female pups drive proliferation independently of TREM2. Indeed, upon ligand activation, CSF1R dimerizes and activates Src kinases; activated Src kinases can either phosphorylate the DAP12 ITAM (for review, [73]) or directly trigger downstream signaling (e.g., PI3K to ERK) to facilitate microglia proliferation. Second, the downregulation of *Pyk2* may partly underlie the profound structural alterations observed in HIV-1 infected microglia [74], as *Pyk2* regulates actin cytoskeleton organization [75].

The complementary use of both statistical and practical significance indexed using *p* values and η_G_^2^, respectively, aids in the interpretation of the present study. In statistics, a *p* value is a conditional probability, whereby its calculation assumes that the null hypothesis (i.e., H_0_) is true; if this probability is less than an arbitrary α value (e.g., 0.05), the null hypothesis is rejected. Although null hypothesis statistical testing is fraught with challenges [76], scientists have relied on these approaches for making inductive inferences for over a century. Herein, consistent with recommendations from the American Psychological Association [77] and the American Statistical Association [78], a standardized alternative inductive approach (i.e., η_G_^2^) was implemented to evaluate the magnitude of the effect. Effect size measures are categorized into two families, including 1) standardized mean differences (e.g., Cohen’s d); and 2) variance explained (e.g., *R*^2^, η_G_^2^). Defined by Olejnik and Algina [40], η_G_^2^ not only provides an estimate of the proportion of variance explained by a predictor but also accounts for the inclusion of a blocking factor (i.e., biological sex) in the experimental design. Reporting and interpreting *p* values and measures of effect size jointly increases our confidence in the transcriptional alterations observed in the present study.

Collectively, primary microglia treated with EcoHIV exhibit a highly specialized and sex-dependent transcriptional state. Independent of biological sex, EcoHIV induced profound microglia hyperproliferation and a stoichiometric imbalance in the TREM2/DAP12 signaling complex. Furthermore, the factor of biological sex regulates precisely how EcoHIV treated microglia respond to the suppression of *Trem2* and compensatory hyperproliferative state. Enhancing our understanding of how HIV-1 alters microglial homeostasis is crucial for the identification of novel therapeutic targets for HAND.

## ACKNOWLEDGEMENTS

We would like to thank Rachel Gray and Dr. Ashlan J. Kunz-Coyne for assistance with EcoHIV plasmid amplification.

## FUNDING

This research was supported in part by grants from NIH (National Institute on Drug Abuse, DA056288; National Institute on Drug Abuse, DA053070) and by the Center of Biomedical Research Excellence (COBRE) in Pharmaceutical Research and Innovation (CPRI, NIH P20-GM130456).

## AUTHOR CONTRIBUTIONS

**Brendan J. Veenstra:** Data curation, Writing-reviewing & editing, Formal analysis. **Kyler J. Nelson:** Data curation, Writing-reviewing & editing. **Taylor R. Elder:** Data curation, Writing-reviewing & editing. **Jill R. Turner:** Writing-reviewing & editing, Funding acquisition. **Kristen A. McLaurin:** Conceptualization, Writing-original draft, Writing-reviewing & editing, Funding acquisition.

## DATA AVAILABILITY STATEMENT

All data is included in this published article and its supplementary information files.

## COMPETING INTERESTS

No author has an actual or perceived conflict of interest with the contents of this article.

## Notes

### Competing Interest Statement

The authors have declared no competing interest.

## REFERENCES

1. UNAIDS. Global HIV & AIDS Statistics. https://www.unaids.org/en/resources/fact-sheet.

2. Autran, B. et al. Positive effects of combined antiretroviral therapy on CD4+ T cell homeostasis and function in advanced HIV disease. Science 277, 112–116 (1997).

3. Koenig, S. et al. Detection of AIDS Virus in Macrophages in Brain Tissue from AIDS Patients with Encephalopathy. Science 233, 1089–1093 (1986).

4. Whitney, J. B. et al. Rapid seeding of the viral reservoir prior to SIV viraemia in rhesus monkeys. Nature 512, 74–77 (2014).

5. Cosenza, M. A., Zhao, M.-L., Si, Q. & Lee, S. C. Human brain parenchymal microglia express CD14 and CD45 and are productively infected by HIV-1 in HIV-1 encephalitis. Brain Pathol 12, 442–455 (2002).

6. Ko, A. et al. Macrophages but not Astrocytes Harbor HIV DNA in the Brains of HIV-1-Infected Aviremic Individuals on Suppressive Antiretroviral Therapy. J Neuroimmune Pharmacol 14, 110–119 (2019).

7. Avalos, C. R. et al. Brain Macrophages in Simian Immunodeficiency Virus-Infected, Antiretroviral-Suppressed Macaques: a Functional Latent Reservoir. mBio 8, e01186–17 (2017).

8. Llewellyn, G. N., Alvarez-Carbonell, D., Chateau, M., Karn, J. & Cannon, P. M. HIV-1 infection of microglial cells in a reconstituted humanized mouse model and identification of compounds that selectively reverse HIV latency. J Neurovirol 24, 192–203 (2018).

9. Li, H., McLaurin, K., Illenberger, J., Mactutus, C. & Booze, R. Microglial HIV-1 Expression: Role in HIV-1 Associated Neurocognitive Disorders. Viruses 13, 924 (2021).

10. Tang, Y. et al. Brain microglia serve as a persistent HIV reservoir despite durable antiretroviral therapy. J Clin Invest 133, e167417 (2023).

11. Nühn, M. M. et al. Microglia Exhibit a Unique Intact HIV Reservoir in Human Postmortem Brain Tissue. Viruses 17, 467 (2025).

12. Wang, Y. et al. Global prevalence and burden of HIV-associated neurocognitive disorder: A meta-analysis. Neurology 95, (2020).

13. Zenebe, Y., Necho, M., Yimam, W. & Akele, B. Worldwide Occurrence of HIV-Associated Neurocognitive Disorders and Its Associated Factors: A Systematic Review and Meta-Analysis. Front. Psychiatry 13, 814362 (2022).

14. Heaton, R. K. et al. HIV-associated neurocognitive disorders before and during the era of combination antiretroviral therapy: differences in rates, nature, and predictors. J Neurovirol 17, 3–16 (2011).

15. Levine, A. J. et al. Multilevel analysis of neuropathogenesis of neurocognitive impairment in HIV. J Neurovirol 22, 431–441 (2016).

16. Rubin, L. H. et al. Microglial activation is inversely associated with cognition in individuals living with HIV on effective antiretroviral therapy. AIDS 32, 1661–1667 (2018).

17. Dos Reis, R. S., et al. Microglial reactivity and nodule formation are associated with Synaptodendritic damage in the brains of people with HIV-1. Brain Pathol 36, e70064 (2026).

18. Di Liberto, G. et al. Neurodegenerative phagocytes mediate synaptic stripping in Neuro-HIV. Brain 145, 2730–2741 (2022).

19. Guha, D., Wagner, M. C. E. & Ayyavoo, V. Human immunodeficiency virus type 1 (HIV-1)-mediated neuroinflammation dysregulates neurogranin and induces synaptodendritic injury. J Neuroinflammation 15, 126 (2018).

20. Campbell, G. R., Rawat, P., To, R. K. & Spector, S. A. HIV-1 Tat Upregulates TREM1 Expression in Human Microglia. J Immunol 211, 429–442 (2023).

21. Jadhav, V. S., Krause, K.-H. & Singh, S. K. HIV-1 Tat C modulates NOX2 and NOX4 expressions through miR-17 in a human microglial cell line. J Neurochem 131, 803–815 (2014).

22. Kong, W. et al. Neuroinflammation generated by HIV-infected microglia promotes dysfunction and death of neurons in human brain organoids. PNAS Nexus 3, pgae179 (2024).

23. Walsh, J. G. et al. Rapid inflammasome activation in microglia contributes to brain disease in HIV/AIDS. Retrovirology 11, 35 (2014).

24. Mamik, M. K. et al. HIV-1 Viral Protein R Activates NLRP3 Inflammasome in Microglia: implications for HIV-1 Associated Neuroinflammation. J Neuroimmune Pharmacol 12, 233–248 (2017).

25. Chivero, E. T. et al. HIV-1 Tat Primes and Activates Microglial NLRP3 Inflammasome-Mediated Neuroinflammation. J Neurosci 37, 3599–3609 (2017).

26. Li, H., McLaurin, K. A., Mactutus, C. F. & Booze, R. M. Microglia proliferation underlies synaptic dysfunction in the prefrontal cortex: implications for the pathogenesis of HIV-1-associated neurocognitive and affective alterations. J Neurovirol 29, 460–471 (2023).

27. Réu, P. et al. The Lifespan and Turnover of Microglia in the Human Brain. Cell Reports 20, 779–784 (2017).

28. Askew, K. et al. Coupled Proliferation and Apoptosis Maintain the Rapid Turnover of Microglia in the Adult Brain. Cell Rep 18, 391–405 (2017).

29. Lawson, L. J., Perry, V. H. & Gordon, S. Turnover of resident microglia in the normal adult mouse brain. Neuroscience 48, 405–415 (1992).

30. Konishi, H. & Kiyama, H. Microglial TREM2/DAP12 Signaling: A Double-Edged Sword in Neural Diseases. Front Cell Neurosci 12, 206 (2018).

31. Hu, B. et al. Insights Into the Role of CSF1R in the Central Nervous System and Neurological Disorders. Front. Aging Neurosci. 13, 789834 (2021).

32. Doorn, K. J. et al. Microglial phenotypes and toll-like receptor 2 in the substantia nigra and hippocampus of incidental Lewy body disease cases and Parkinson’s disease patients. Acta Neuropathol Commun 2, 90 (2014).

33. Villacampa, N. et al. Proliferating Microglia Exhibit Unique Transcriptional and Functional Alterations in Alzheimer’s Disease. ASN Neuro 17, 2506406 (2025).

34. Potash, M. J. et al. A mouse model for study of systemic HIV-1 infection, antiviral immune responses, and neuroinvasiveness. Proc Natl Acad Sci U S A 102, 3760–3765 (2005).

35. Kim, B.-H. et al. EcoHIV Infection of Primary Murine Brain Cell Cultures to Model HIV Replication and Neuropathogenesis. Viruses 16, 693 (2024).

36. Bobotis, B. C. et al. Sex differences of microglia in the healthy brain from embryonic development to adulthood and across lifestyle influences. Brain Res Bull 202, 110752 (2023).

37. Alfar, H. R. et al. Protocol for optimizing production and quality control of infective EcoHIV virions. STAR Protoc 4, 102368 (2023).

38. Livak, K. J. & Schmittgen, T. D. Analysis of relative gene expression data using real-time quantitative PCR and the 2(-Delta Delta C(T)) Method. Methods 25, 402–408 (2001).

39. Schneider, C. A., Rasband, W. S. & Eliceiri, K. W. NIH Image to ImageJ: 25 years of image analysis. Nat Methods 9, 671–675 (2012).

40. Olejnik, S. & Algina, J. Generalized eta and omega squared statistics: measures of effect size for some common research designs. Psychol Methods 8, 434–447 (2003).

41. Cohen, J. Statistical Power Analysis for the Behavioral Sciences. Academic Press (1969).

42. Streit, W. J., Graeber, M. B. & Kreutzberg, G. W. Functional plasticity of microglia: a review. Glia 1, 301–307 (1988).

43. Davalos, D. et al. ATP mediates rapid microglial response to local brain injury in vivo. Nat Neurosci 8, 752–758 (2005).

44. Nimmerjahn, A., Kirchhoff, F. & Helmchen, F. Resting microglial cells are highly dynamic surveillants of brain parenchyma in vivo. Science 308, 1314–1318 (2005).

45. Sierra, A., Tremblay, M.-È. & Wake, H. Never-resting microglia: physiological roles in the healthy brain and pathological implications. Front Cell Neurosci 8, 240 (2014).

46. Colton, C. A. et al. Expression profiles for macrophage alternative activation genes in AD and in mouse models of AD. J Neuroinflammation 3, 27 (2006).

47. Ransohoff, R. M. A polarizing question: do M1 and M2 microglia exist? Nat Neurosci 19, 987–991 (2016).

48. Lee, S. C., Liu, W., Brosnan, C. F. & Dickson, D. W. GM-CSF promotes proliferation of human fetal and adult microglia in primary cultures. Glia 12, 309–318 (1994).

49. Gómez-Nicola, D., Fransen, N. L., Suzzi, S. & Perry, V. H. Regulation of microglial proliferation during chronic neurodegeneration. J Neurosci 33, 2481–2493 (2013).

50. McDonough, A. et al. Ischemic preconditioning induces cortical microglial proliferation and a transcriptomic program of robust cell cycle activation. Glia 68, 76–94 (2020).

51. Kurki, P., Vanderlaan, M., Dolbeare, F., Gray, J. & Tan, E. M. Expression of proliferating cell nuclear antigen (PCNA)/cyclin during the cell cycle. Exp Cell Res 166, 209–219 (1986).

52. Bravo, R. & Macdonald-Bravo, H. Existence of two populations of cyclin/proliferating cell nuclear antigen during the cell cycle: association with DNA replication sites. J Cell Biol 105, 1549–1554 (1987).

53. Kelman, Z. PCNA: structure, functions and interactions. Oncogene 14, 629–640 (1997).

54. Cuylen, S. et al. Ki-67 acts as a biological surfactant to disperse mitotic chromosomes. Nature 535, 308–312 (2016).

55. Gerdes, J., Schwab, U., Lemke, H. & Stein, H. Production of a mouse monoclonal antibody reactive with a human nuclear antigen associated with cell proliferation. Int J Cancer 31, 13–20 (1983).

56. Gerdes, J. et al. Cell cycle analysis of a cell proliferation-associated human nuclear antigen defined by the monoclonal antibody Ki-67. J Immunol 133, 1710–1715 (1984).

57. Cosenza-Nashat, M. et al. Human immunodeficiency virus infection inhibits granulocyte-macrophage colony-stimulating factor-induced microglial proliferation. J Neurovirol 13, 536–548 (2007).

58. Filipowicz, A. R. et al. Proliferation of Perivascular Macrophages Contributes to the Development of Encephalitic Lesions in HIV-Infected Humans and in SIV-Infected Macaques. Sci Rep 6, 32900 (2016).

59. Avalos, B. et al. Cannabis Use and Cannabidiol Modulate HIV-Induced Alterations in TREM2 Expression: Implications for Age-Related Neuropathogenesis. Viruses 16, 1509 (2024).

60. Fields, J. A. et al. Alterations in brain TREM2 and Amyloid-β levels are associated with neurocognitive impairment in HIV-infected persons on antiretroviral therapy. J Neurochem 147, 784–802 (2018).

61. Yao, H. et al. Distinct Signaling Pathways Regulate TREM2 Phagocytic and NFκB Antagonistic Activities. Front Cell Neurosci 13, 457 (2019).

62. Zhao, Y. et al. TREM2 Is a Receptor for β-Amyloid that Mediates Microglial Function. Neuron 97, 1023–1031.e7 (2018).

63. Ruganzu, J. B. et al. Downregulation of TREM2 expression exacerbates neuroinflammatory responses through TLR4-mediated MAPK signaling pathway in a transgenic mouse model of Alzheimer’s disease. Mol Immunol 142, 22–36 (2022).

64. Achim, C. L. et al. Increased accumulation of intraneuronal amyloid beta in HIV-infected patients. J Neuroimmune Pharmacol 4, 190–199 (2009).

65. Li, H. et al. Intraneuronal β-Amyloid Accumulation: Aging HIV-1 Human and HIV-1 Transgenic Rat Brain. Viruses 14, 1268 (2022).

66. McLaurin, K. A., Li, H., Ritchie, A., Booze, R. M. & Mactutus, C. F. IMPAIRED BRIDGING OF TEMPORAL DISCONTINUITIES IN OLDER ADULT HIV-1 TG RATS. Preprint at 10.64898/2026.04.06.716768 (2026).

67. Gisslén, M. et al. CSF concentrations of soluble TREM2 as a marker of microglial activation in HIV-1 infection. Neurol Neuroimmunol Neuroinflamm 6, e512 (2019).

68. Farias Quipildor, G. E., et al. Differential downstream signaling in microglia lacking Alzheimer’s-related TREM2 or its adaptor TYROBP/DAP12. Mol Neurodegener Adv 2, 8 (2026).

69. Bouchon, A., Dietrich, J. & Colonna, M. Cutting edge: inflammatory responses can be triggered by TREM-1, a novel receptor expressed on neutrophils and monocytes. J Immunol 164, 4991–4995 (2000).

70. Chung, I.-C. et al. Pyk2 activates the NLRP3 inflammasome by directly phosphorylating ASC and contributes to inflammasome-dependent peritonitis. Sci Rep 6, 36214 (2016).

71. Knight, A. C., Brill, S. A., Queen, S. E., Tarwater, P. M. & Mankowski, J. L. Increased Microglial CSF1R Expression in the SIV/Macaque Model of HIV CNS Disease. J Neuropathol Exp Neurol 77, 199–206 (2018).

72. Irons, D. L., Meinhardt, T., Allers, C., Kuroda, M. J. & Kim, W.-K. Overexpression and activation of colony-stimulating factor 1 receptor in the SIV/macaque model of HIV infection and neuroHIV. Brain Pathol 29, 826–836 (2019).

73. Lee, J.-W., Lee, I.-H., Iimura, T. & Kong, S. W. Two macrophages, osteoclasts and microglia: from development to pleiotropy. Bone Res 9, 11 (2021).

74. Renner, N. A. et al. Microglia activation by SIV-infected macrophages: alterations in morphology and cytokine secretion. J Neurovirol 18, 213–221 (2012).

75. Du, Q. S. et al. Inhibition of PYK2-induced actin cytoskeleton reorganization, PYK2 autophosphorylation and focal adhesion targeting by FAK. J Cell Sci 114, 2977–2987 (2001).

76. Nuzzo, R. Scientific method: Statistical errors. Nature 506, 150–152 (2014).

77. Wilkinson, L. & Task Force on Statistical Inference. Statistical methods in psychology journals: Guidelines and explanations. American Psychologist 54, 594–604 (1999).

78. Wasserstein, R. L. & Lazar, N. A. The ASA Statement on *p* -Values: Context, Process, and Purpose. The American Statistician 70, 129–133 (2016).

