## Supplementary Information for "EcoHIV Treated Primary Microglia Exhibit a Highly Specialized and Sex-Dependent Transcriptional State"

Lexington, KY 40508

**Number of Tables, Figures**: 1, 5

**Address proofs and correspondence to:**

Kristen A. McLaurin, Ph.D.

Assistant Professor

Department of Pharmaceutical Sciences

College of Pharmacy

789 South Limestone Street

University of Kentucky

Lexington, KY 40508

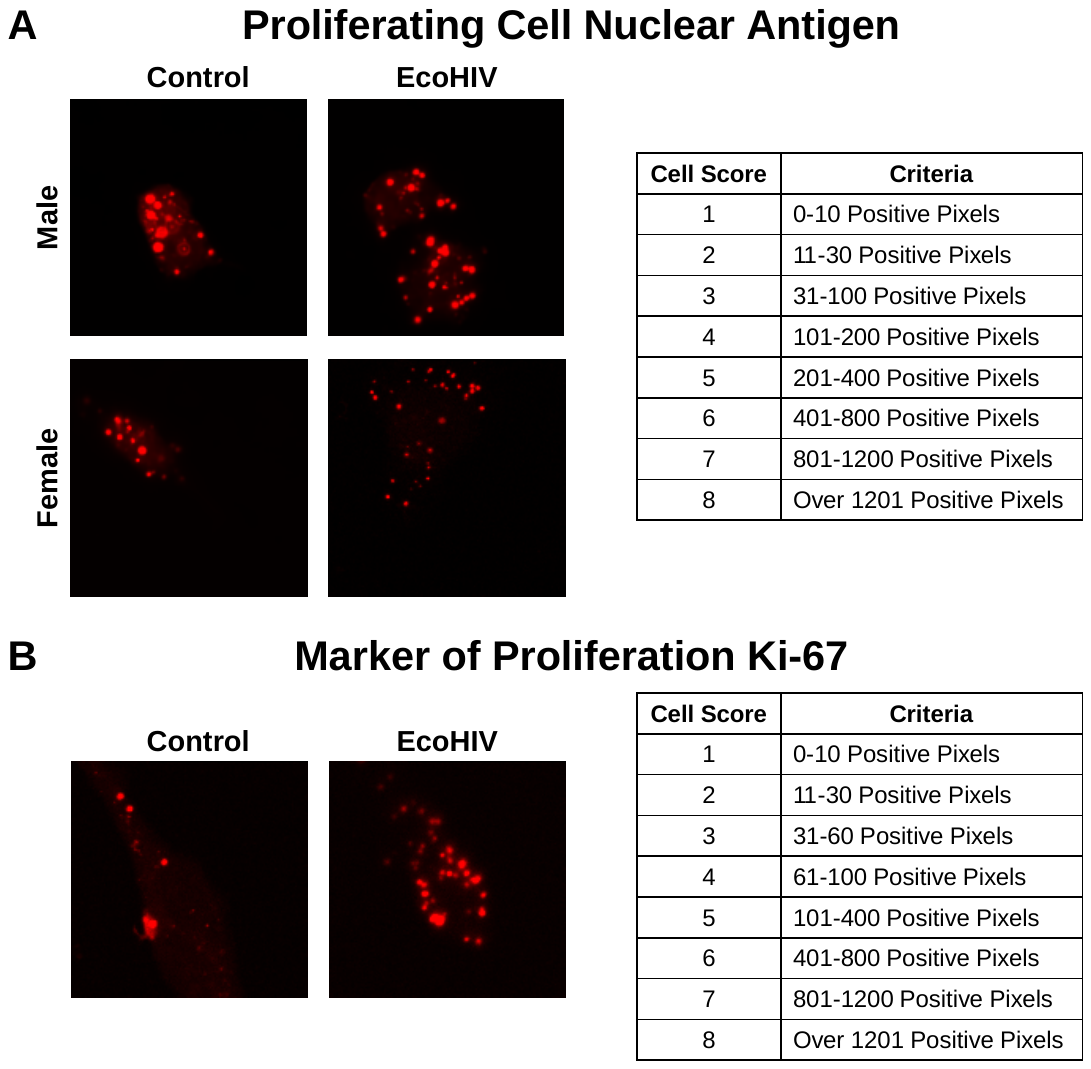

**Supplementary Figure 1.** Representative images and criteria for the quantitative analysis of *in situ* hybridization for (**A**) proliferating cell nuclear antigen (*Pcna*) and (**B**) marker of proliferation Ki-67 (*Mki67*).

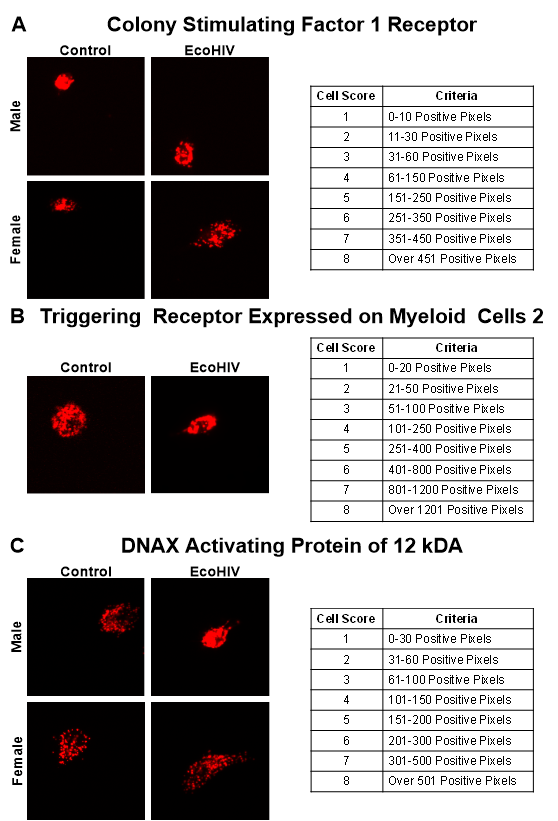

**Supplementary Figure 2.** Representative images and criteria for the quantitative analysis of *in situ* hybridization for (**A**) colony stimulating factor 1 receptor (*Csf1r*), (**B**) triggering receptor expressed on myeloid cells 2 (*Trem2*) and (**C**) DNAX activating protein of 12 kDa (*Dap12*).
